# N-terminal truncation diversifies USP18 function during early myogenesis

**DOI:** 10.64898/2026.10.12.758511

**Authors:** Tom I. Nijssen, Simeon D. Draganov, Milad Shademan, Sarah Flannery, Iolanda Vendrell, Benedikt M. Kessler, Adan Pinto-Fernandez, Vered Raz

## Abstract

Ubiquitin-specific protease 18 (USP18) is primarily studied for its dual role in type-I interferon (IFN-I) signaling, acting as an ISG15 deISGylase and as a non-catalytic inhibitor of IFNAR signaling. At basal levels, however, USP18 also regulates muscle cell development, independently of IFN-I signaling. USP18 exists as a canonical long isoform (LF) and a short isoform (SF) generated by N-terminal truncation. Endogenous SF accumulates in the nucleus during muscle differentiation, but its cellular functions remain poorly understood. Here, we compared USP18 SF and LF in muscle cells stably overexpressing each isoform. We show that their distinct N-termini are associated with marked differences in protein stability, subcellular localization, and protein–protein interactions. SF displayed a substantially expanded interactome, with increased association with nuclear proteins, chaperones, and regulators of protein homeostasis. Functionally, SF enhanced an early stage of myogenic differentiation and was associated with changes in alternative splicing, including key sarcomere genes. Together, these findings identify the SF USP18 proteoform as a potential regulator of muscle differentiation and highlight the N-terminus as a determinant of non-canonical USP18 functions and proteoform diversity.

**Graphical Abstract:** N-terminal truncation alters USP18 stability, localization, and protein interactions, potentially enabling a distinct function in early myogenic differentiation.

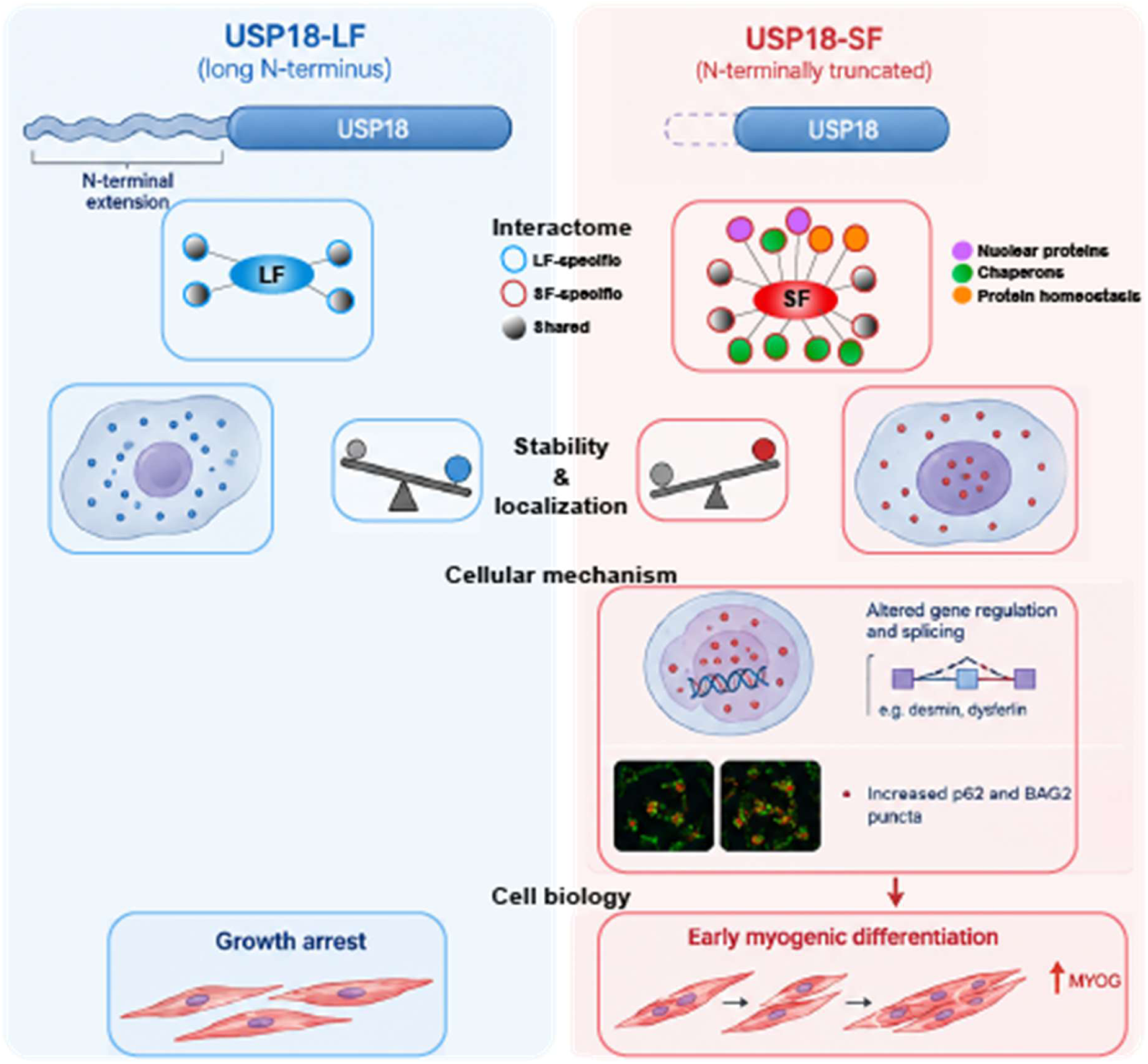

## Introduction

Ubiquitin-Specific Peptidase 18 (USP18) is a key negative regulator of the type-I interferon (IFN-I) stimulated gene (ISG) response and the primary deISGylating enzyme that cleaves the ubiquitin-like protein interferon-stimulated gene 15 (ISG15) from substrate proteins. By removing ISG15 from target proteins, USP18 negatively regulates the ISGylation pathway and limits excessive or prolonged IFN-I responses (Honke et al., 2016; Kang & Jeon, 2020). ISG15 expression is rapidly induced by IFN-I and plays a central role in antiviral and innate immune responses (Perng & Lenschow, 2018). In addition to its catalytic activity, USP18 directly attenuates IFN-I signaling by interacting with STAT2 and the interferon receptor subunit IFNAR2, a non-catalytic/scaffolding function (Benavente-Diaz et al., 2021; Faralli & Dilworth, 2012; Yin et al., 2023). This prevents productive receptor complex formation and limits downstream JAK-STAT signaling as a negative feedback loop (Arimoto et al., 2017). USP18 can also interact with E2-E3 ligase complexes and co-regulate ubiquitination in diverse cancers (Hou et al., 2021; Napoletano et al., 2026), suggesting additional potential scaffold functions for USP18 that may depend on cell-type context.

Recently, we showed that USP18 alters muscle cell differentiation (Olie et al., 2023). Notably, these effects occur independently of IFN-I and its deISGylating activity (Nijssen et al., 2026), pointing towards another evolutionary conserved non-canonical role for USP18 in myogenesis. Myogenesis is tightly regulated by the sequential expression of myogenic regulatory factors (MRFs), amongst which MYOG (myogenin) induces the expression of cell-cycle exit regulators, myoblast fusion, and development of the contractile apparatus (Benavente-Diaz et al., 2021; Faralli & Dilworth, 2012; Yin et al., 2023). In vitro, myogenesis of cultured myoblasts is initiated by growth arrest following serum starvation, followed by myoblast fusion and myofiber maturation. This process can be monitored by the transient induction of MYOG, followed by expression of MyHC as cells progress toward mature myofibers (Chal & Pourquié, 2017). Although transcriptional regulation is essential for successful myogenic differentiation, the process also depends on protein homeostasis, which provides rapid and reversible control of protein function, stability, localization and protein-protein interactions (Zan et al., 2025). How USP18 regulates myogenesis is currently not fully understood.

USP18 exists as two protein isoforms that arise from alternative translation initiation sites within the USP18 mRNA. The long isoform (LF) is translated from an upstream non-canonical CUG start codon, whereas the short isoform (SF) is initiated from a downstream AUG codon, resulting in an N-terminally truncated protein lacking the first 35 amino acids (Burkart et al., 2012). Both isoforms retain the catalytic triad required for deISGylation and the domains involved in interferon signaling but differ in their subcellular localization. The LF is predominantly cytoplasmic, whereas the SF displays enhanced nuclear localization (Arimoto et al., 2023; Burkart et al., 2012). We previously reported the accumulation of USP18 isoforms during muscle cell differentiation, with the short isoform showing a particularly pronounced increase in nuclear fraction. (Olie et al., 2023), raising the hypothesis that nuclear USP18 may contribute to a non-canonical function during myogenesis.

In this study, we investigated the isoform-specific roles of USP18 during muscle cell differentiation. We hypothesized that the short USP18 isoform may promote myogenesis by regulating nuclear proteostasis through its scaffold function.

## Methods

### USP18 expression vectors

The USP18-LF isoform was made by conversion of Methionine-36 to Alanine using the primer set 1 as described in (Nijssen et al., 2026). The USP18-SF isoform was made by deletion of 105 nucleotides from the 5’ end of the gene, deleting 35 amino acids from the N-terminus with the primer set 2. List of primers can be found in **Table S1**. Mutagenesis was performed by in-vitro assembly according to (García-Nafría et al., 2016).

### Cell Culture

Immortalized human muscle cells (home-made clone 2417 (Mallon et al., 2025)), were used in this study and are different from the cells previously described for the USP18 knockdown experiments (Olie et al., 2023). Cells were transduced with lentivirus particles as detailed in (Raz et al., 2006). Subsequently, transduction efficiency was assessed using GFP. The human muscle cells were cultured in growth medium consisting of Gibco F10 medium, supplemented with 15% fetal calf serum, 1% penicillin/streptomycin, 0.4ug/ml dexamethasone (Sigma-Aldrich), 1ng/ml basic fibroblast growth factor. For differentiation, muscle cells medium consisting of DMEM supplemented with 2% horse serum was used.

Experiments in proliferation conditions were conducted at approximately 70% confluence, whereas differentiation experiments were performed at confluence approximating 100% for 1, 2 or 3 days. For protein stability assay, cells were incubated with 20 ug/ml cycloheximide for the mentioned times.

### Cell lysis, subcellular fractionation, Western blotting, and immunoprecipitation

For bulk protein assay cells were lysed on ice using RIPA (HEPES pH7.0 10mM, NaCl 150mM, MgCl 5mM, Triton 1%, Glycerol 10%, SDS 2%) freshly supplemented with protease inhibitor (Roche) and DTT 1mM, and subsequently sonicated. Supernatant was mixed with Laemmli sample buffer (Biorad) with 250 mM DTT and boiled at 95 degrees Celsius for 10 minutes. Equal amounts of protein were separated on a 10% polyacrylamide gel and subsequently blotted onto a PVDF membrane for 1 hour at 320 mA at 4 degrees Celsius. The membrane was blocked using PBS with 0.1% Tween (PBST) with 5% milk for a minimum of 30 minutes at room temperature. Primary antibodies were incubated overnight at 4 degrees Celsius and after washing incubated with secondary antibodies for 30 minutes at room temperature. Blots were imaged with Odyssey CLx (LI-COR). Primary and secondary antibodies are listed in **Table S2**.

For the sub-cellular fractionation assay, cells were lysed using the cytosolic buffer (HEPES pH8 50mM, NaCl 150mM) freshly supplemented with protease inhibitor (Roche) and DTT 1mM on ice for 30 minutes. The cytoplasmic fraction was separated from the nuclear fraction by centrifugation at 3500xg at 4 degrees Celsius for 10 minutes. The pellet was subsequently washed with PBS, and nuclear proteins were lysed with the nuclear lysis buffer (HEPES pH8 50mM, NaCl 150mM, Sodium deoxycholate 0.1% wv, Triton 0.1%) freshly supplemented with protease inhibitor (Roche) and DTT 1mM on ice for 10 minutes. The nuclear proteins were separated from the insoluble proteins by centrifugation at 7800xg for 10 minutes at 4°C. The supernatant containing the nuclear proteins was collected to a new tube and the pellet was discarded. The cytoplasmic and nuclear fractions were subsequently used for Western blot analysis.

Immunoprecipitation was performed on cells in proliferation conditions and after 1 day of differentiation. Cells were lysed using IP buffer (20 mM HEPES pH 8, 150 mM NaCl, 0.2% NP-40, 10 % glycerol, 5 mM NEM) freshly supplemented with protease inhibitor (cOmplete^™^, Mini, EDTA-free Protease Inhibitor Cocktail, Sigma Aldrich). For LC MS/MS lysates were incubated with anti-Flag (M2, Sigma) antibodies or without antibodies (beads only) at 4°C overnight, followed by a one-hour incubation with protein G Sepharose beads slurry (Invitrogen, #15920-10) for one hour at 4°C. For Co-IP validation of the mass spectrometry analysis was made with FLAG-Trap (Chromotek), according to the manufacturer’s protocol.

### Proteomics analysis (sample preparation, LC MS/MS and Data analysis)

Co-IP elution was made using 2× Gel Loading buffer (GLB) with 25 mM DTT in two steps: first, using 50 µL of GLB and boiled at 90 °C for 5 min, next using 60 µL of GLB and boiled at 90 °C for 10 min. Finally, both the elution sets were combined. 10% of the eluate was used for Western blot analysis and the remainder for mass spectrometry. Eluate samples were reduced using 0.8 µL of 1.25 M DTT per 100 µL (final concentration of 10 mM DTT per sample). Alkylation of the disulfides was done using 10 µL of 200 mM Iodoacetamide (IAA) per 100 µL of sample (final concentration of 20 mM). The sample then was acidified using 10 µL of 1.1% phosphoric acid (final pH of sample ≤ 1) and washed with 620 µL of binding/wash buffer (90% Methanol, 100 mM TEAB, pH 7.55). Samples were next loaded onto S-trap (Protifi) columns and centrifuged at 4000 × *g* for 30 s to bind to the column. The columns were washed with 400 µL of the same buffer 4 times by centrifuging at 4000 × *g* for 30 s each. The residual buffer was removed by final centrifugation at 4000 × *g* for 1 min. Finally, the filters were transferred to new 2 mL collection tubes, and 125 µL of Trypsin (1:10 wt:wt ratio) was added to the filter. Tubes were kept at room temperature without shaking for overnight digestion. Peptides were eluted from the S-trap columns using Buffer 1 (50 mM TEAB in water, pH 8.5), Buffer 2 (0.2% formic acid in water), and Buffer 3 (50% acetonitrile in water) successively. Each time, 80 µL of the buffer was applied to the column and centrifuged at 4000 × *g* for 1 min. Eluted proteins were dried using a vacuum evaporator (SP Genevac miVac centrifugal evaporator miVacATS life sciences scientific products). Dried samples were stored at −80 °C. They were resuspended prior MS analysis in mass spectrometry grade 98% water + 2% acetonitrile + 0.1% TFA to a total volume of 15 µL.

Peptides were analyzed by LC-MS/MS using an Evosep ONE LC system connected to the Tims-TOF-Pro from Bruker. Tryptic peptides were loaded onto Evotip Pure (EV2011) following manufacturer protocol. All centrifugation steps were performed at 800 x g for 60 seconds. Then, peptides were chromatographically separated on a 15 cm x 150 µm x 1.5 µm analytical column (EV1137, Evosep) using the 30 samples per day (spd) gradient method of an Evosep ONE LC system. The Tims-TOF-Pro was operated in dia-PASEF mode using 16 dia-PASEF scans per TIMS-MS scan. The ion mobility range was set to 0.6 – 1.6 Vs/cm2. Each mass window isolated was 26 m/z wide, with a mass overlap of 1Da per mobility window, ranging from 400-1200 m/z with an ion mobility-dependent collision energy that increased linearly from 20 eV to 59 eV between 0.6 – 1.6 Vs/cm2.

Mass spectrometry data were analyzed using DIA-NN (version 1.9.2) in library-free mode against the UniProtKB human proteome database (downloaded in November 2023; 20,426 sequences), supplemented with common contaminants. Data were searched using default settings, with a 1% false discovery rate (FDR) applied at the precursor level and match-between-runs (MBR) enabled. Trypsin was specified as the proteolytic enzyme, allowing up to one missed cleavage. Carbamidomethylation of cysteine was specified as a fixed modification, while methionine oxidation was included as a variable modification. N-terminal methionine excision was enabled.

Proteome analysis, including the identification of significantly enriched binders in FLAG immunoprecipitation (FLAG-IP) samples compared with beads-only controls using paired analysis, was performed in Perseus software (version 2.0.11). Significant interactors were determined with FDR 5%. Lists of LF and SP significant interactors per cell condition are in **Table S3A.** Major sources of variation between samples were assessed by principal component analysis (PCA). From this pool of significantly enriched binders (5% FDR), isoform-specific binders were subsequently identified using a t-test (p < 0.05), with protein abundance normalized to the immunoglobulin (Ig) levels in each IP sample. A list of SP-specific interactors is in **Table S3B.**

### RNA extraction, qPCR, library preparation, RNA sequencing, and alternative slicing analysis

Total RNA was isolated from cultured cells using the RNeasy Mini Kit (QIAGEN) according to the manufacturer’s instructions, followed by an on-column RNase-free DNase treatment (QIAGEN) to remove residual genomic DNA. RNA was stored at -80 °C. RNA concentration was quantified using Qubit (ThermoFisher). For cDNA synthesis, 500 ng of RNA was reverse transcribed using the QuantiTect Reverse Transcription Kit (Qiagen, #205311) with random hexamer primers according to the manufacturer’s protocol. Quantitative PCR (qPCR) was performed using the QuantiNova SYBR Green PCR Kit (Qiagen, #208057). Three nanograms of cDNA were used per reaction, and all samples were analyzed in technical triplicate. Gene expression levels were determined using primers specific to USP18 and the indicated housekeeping gene(s). Primer sequences are listed in **Table S1**.

For RNA sequencing RNA integrity was checked on an RNA 6000 Nano Agilent Lab-on-a-Chip kit with the Bioanalyzer System before cDNA library preparation. RIN score >8 and 100 ng total RNA per sample was used for poly(A) library preparation using 7 PCR amplification cycles. Library size distribution was verified on an Agilent Lab-on-a-Chip and quantified using Qubit prior to pooling. Illumina paired-end sequencing was performed on a NovaSeq 6000 system using 150-bp paired-end reads and v1.5 chemistry. All reads in FASTQ format were first filtered using Cutadapt (v2.10) to remove all remaining adapter sequences, and quality control (QC) of the FASTQ files was assessed with the MultiQC program in Python. The remaining reads were aligned to the Ensembl transcriptome version 104 using STAR (v2.7.5a), including UMI-based deduplication with UMI-Tools (v1.1.1), generating a transcriptome-based alignment in BAM format. The same Ensembl transcript annotation (version 104) was used to create a customized transcript annotation GTF file for all Ensembl-annotated transcripts.

Reads were quantified using featureCounts (v2.0.1). Significant differentially expressed genes were selected with an eBayes *t*-test with FDR < 0.05, and genes were assigned to the LF or SF according to the log_2_ fold change direction. A list of dysregulated genes is in **Table S4**. All analyses were performed using RStudio Software (RStudio 2022.02.3, Build 492) with R Statistical Software (v4.2.3). Volcano plot visualization of differential expression was made with ggVolcanoR (https://ggvolcanor.erc.monash.edu/).

Alternative splicing was quantified and analyzed with rMATS-turbo (https://github.com/Xinglab/rmats-turbo). rMATS was run with the Ensembl GRCh38.115 annotation and the following parameters: --novelSS (novel splice-site detection enabled), -t paired, --libType fr-unstranded, --readLength 150, --nthread 4 and --drop-zero-read-replicates-for-stat, which excludes replicates with zero reads from the statistical model. Analyses were performed on the JCEC output (junction counts plus reads on the target exon) for all five event classes: skipped exon (SE), mutually exclusive exons (MXE), alternative 5′ splice site (A5SS), alternative 3′ splice site (A3SS) and retained intron (RI). Duplicate-marked BAM files for LF (n = 3) and SF (n = 3) were supplied as Level 1 and Level 2 biological groups, respectively. Therefore, ΔPSI represents the difference between the mean exon inclusion levels of the two groups, PSI(LF) − PSI(SF). A positive ΔPSI corresponds to higher exon inclusion in LF. For each event, rMATS reports the PSI per replicate in each group as:

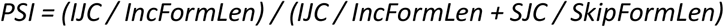

Where IJC and SJC are the inclusion and skipping junction, read counts and IncFormLen and SkipFormLen are the effective lengths of the two isoforms.

Filtering was applied in three steps. First, events with a missing inclusion-level value (NA) in any replicate of either group (IncLevel1 or IncLevel2), second, events with FDR < 0.05 were selected. Third, an effect-size threshold of |ΔPSI| ≥ 0.1 was applied, and events were assigned to the LF direction (ΔPSI ≥ +0.1) or the SF direction (ΔPSI ≤ −0.1); events with |ΔPSI| < 0.1 were discarded. Significant differences between LF and SF were tested for each event class using a two-sided exact binomial test. P values were adjusted using the Benjamini-Hochberg procedure, and the overall LF:SF distribution was assessed with a chi-square test. A list of significant alternative splicing transcripts is in **Table S5.** Individual events were inspected in the Integrative Genomics Viewer (IGV) v2.19.5 using the duplicate-marked BAM files. Sashimi plots were generated in IGV per group with junction read counts displayed on the arcs.

### Cell growth assay

Cell proliferation was assessed on a 96-well plate using the IncuCyte (version 2020B, Sartorius). Live-cell imaging was performed every 3 hours while growing at 37 degrees Celsius with 5% CO2. The growth rate was calculated after normalization to t=0.

### Immunofluorescence, Imaging, and Image quantification

Immunofluorescence staining was performed on a 96-well plate using cultured muscle cells. Cells were fixed with 4% formaldehyde in PBS, followed by permeabilization with PBS with 1% Triton X-100. Cells were incubated with primary antibodies for 1 hour at room temperature, followed by 30 minutes of secondary antibodies in combination with DAPI (1:1000) after washing. Primary and secondary antibodies can be found in **Table S2**. Imaging and image quantification were made with CellInsight CX7 microscope and the built-in software (Thermo Fisher). Nuclear localization and fusion index, and the overlap between MYOG and USP18 were measured with the co-localization toolbox. Autophagy puncta were segmented and measured with the Spot detector toolbox.

### Statistical analysis and online bioinformatics

Statistical analysis and plots were made with GraphPad Prism (version 10.2.3). Statistical significance was determined by one-way or two-way ANOVA with Sidak correction.

Gene set or protein enrichment was made in STRING bioinformatics (https://string-db.org/), DAVID (https://davidbioinformatics.nih.gov/) using our gene set as background, or with ShinyGO v0.85 (https://bioinformatics.sdstate.edu/go/) cut-off of 0.05 FDR. Non-redundant terms were selected and sorted by fold enrichment.

Models of USP18 folding were predicted using Alphafold3. Functional domains were annotated based on UniProt referencing (Arimoto et al., 2017). Visualization of prediction and domains was made in Pymol (version 3.1.6.1). Degron prediction was made at degronopedia.com.

## Results

### USP18 isoforms exhibit distinct localization, structural features, and functions during myogenic differentiation

To investigate USP18 isoform-specific effects on USP18 muscle cell biology, we stably overexpressed either the long (LF) or short (SF) USP18 isoform in immortalized human muscle cells. The LF was made by silencing the alternative start codon M35 using the M36A mutation (**Figure 1A**). The SF was made by deleting the first 35 amino acids (**Figure 1A**). Both isoforms were N-terminally tagged with FLAG and cloned into a lentiviral IRES-GFP vector, with GFP serving as a marker to select and monitor stable cell lines (**Figure 1A**). Overexpression of both USP18 isoforms was confirmed by qPCR, which demonstrated comparable transcript levels, while endogenous USP18 transcripts were undetectable (**Figure 1B**). Western blot analysis verified protein expression of both isoforms, with SF expressed at higher levels than LF and migrating approximately 4 kDa lower, consistent with the predicted molecular weight difference (**Figure 1C**). Under proliferation conditions, endogenous USP18 protein was not detected (**Figure 1C**). Immunofluorescence showed that LF localized predominantly to cytoplasm, whereas SF showed marked nuclear enrichment (**Figure 1D**). Consistent with this observation, nuclear FLAG mean fluorescence intensity (MFI) was nearly threefold higher in SF-expressing cells than in LF-expressing cells (**Figure 1E**). We further quantified subcellular distribution by calculating the nuclear-to-cytoplasmic fluorescence ratio. LF-expressing cells exhibited a ratio of 0.5, indicating predominantly cytoplasmic localization, whereas SF-expressing cells exhibited a ratio of 3.3, consistent with preferential nuclear accumulation (**Figure 1E**). Subcellular fractionation followed by Western blot analysis confirmed the enrichment of SF in the nuclear fraction (**Figure 1F**). Together, these findings demonstrate that in muscle cells LF and SF showed opposite subcellular localization.

**Figure 1.**
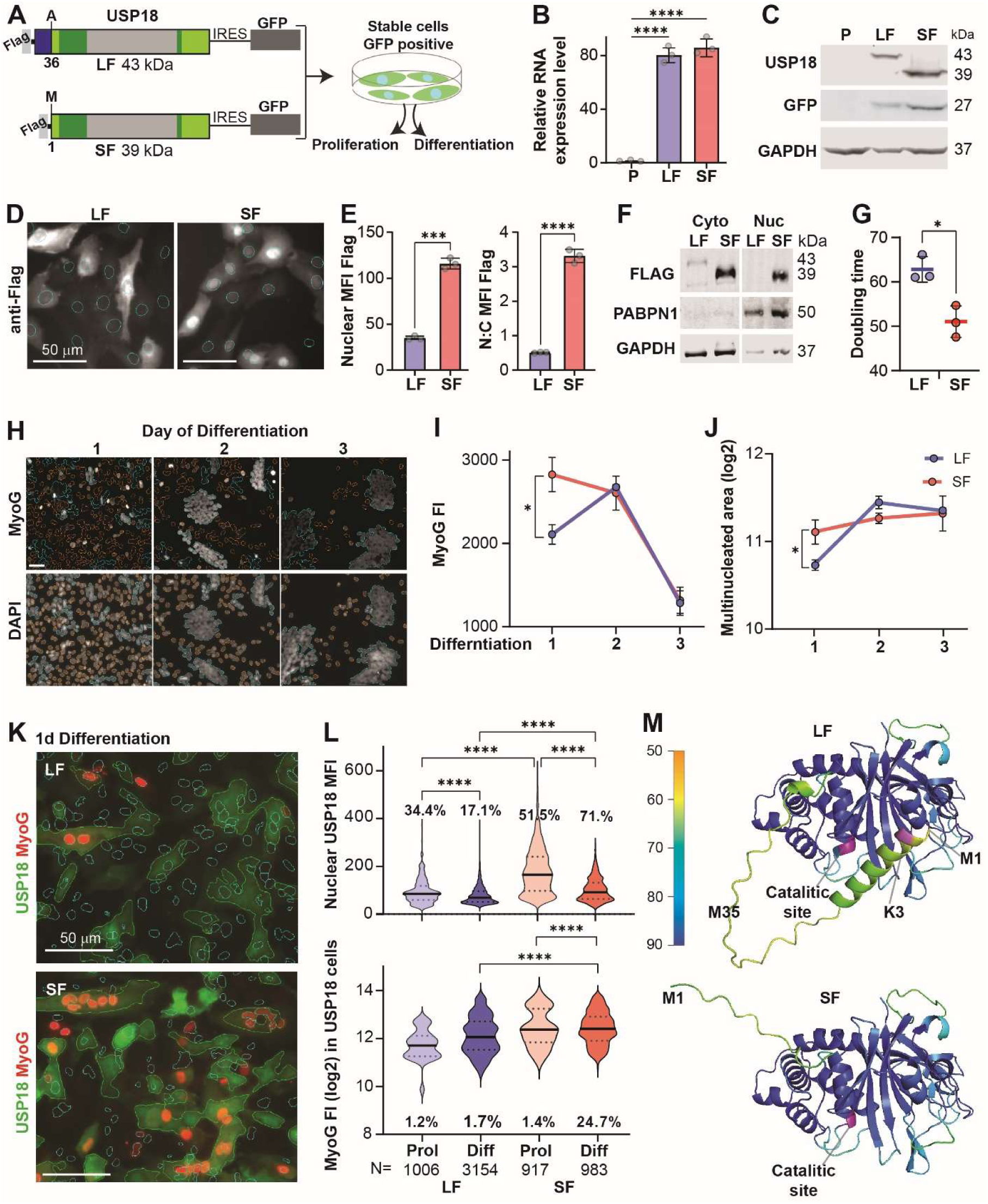
USP18-SF preferentially localizes the nucleus and promotes early myogenic differentiation. **A.** Schematic of the lentiviral constructs used to generate stable human muscle cell lines expressing FLAG-tagged USP18 long form (LF, 43 kDa) or short form (SF, 39 kDa). In the LF, the alternative translation initiation site (M36) was mutated (M36A). SF was generated by deleting the first 35 amino acids. In blue, the N-terminus unique to LF, in green the binding domains of the IFNAR (dark green) and STAT (light green). Both constructs were cloned into an IRES-GFP vector. **B.** Quantitative PCR analysis confirming comparable overexpression of LF and SF transcripts relative to parental (P) cells. Data are presented as mean ± SE. \*\*\*\**P* < 0.0001. **C.** Western blot analysis confirming expression of LF and SF proteins. GFP expression verified transgene expression and GAPDH served as the loading control. **D.** Representative immunofluorescence images of FLAG staining in LF- and SF-expressing cells. Nuclei are outlined for quantification. Scale bar, 50 μm. **E.** Quantification of nuclear FLAG mean fluorescence intensity (MFI) and the nuclear-to-cytoplasmic (N:C) fluorescence ratio, demonstrating significantly greater nuclear accumulation of USP18-SF. Data are shown as mean ± SE. \*\*\**P* < 0.001; \*\*\*\**P* < 0.0001. **F.** Western blot analysis of cytoplasmic (Cyto) and nuclear (Nuc) fractions confirming preferential nuclear localization of SF. PABPN1 and GAPDH served as nuclear and cytoplasmic fraction controls, respectively. **G.** Cell doubling time of LF- and SF-expressing cells during proliferation. Data are presented as mean ± SE. \**P* < 0.05. **H.** Representative immunofluorescence images of MYOG expression during 1, 2, and 3 days of differentiation. MYOG-positive cells are shown together with DAPI nuclear staining. Scale bar, 100 μm. **I.** Quantification of MYOG fluorescence intensity during differentiation. Data are presented as mean ± SE. *P* < 0.05. **J.** Quantification of multinucleated cell areas during differentiation. Data are shown as mean ± SE. \**P* < 0.05. **K.** Representative immunofluorescence images of USP18 (green) and MYOG (red) after 1 day of differentiation in LF- and SF-expressing cells. Scale bar, 50 μm. **L.** Single-cell quantification of nuclear USP18 fluorescence intensity (top) and MYOG fluorescence intensity in USP18-positive cells (bottom) during proliferation (Prol) and after 1 day of differentiation (Diff). Percentages indicate the proportion of USP18-positive nuclei (top) and MYOG-positive nuclei (bottom). Violin plots show the median and quartiles. \*\*\*\**P* < 0.0001. Violin plots are from >1000 USP18-positive cells. **M.** Folding predictions of USP18-LF and USP18-SF. Color scale indicates the prediction of confidence (pLDDT), from 50 to 90 percentage. The first methionine (M1) and alternative methionine (M35) are indicated. The ubiquitinated lysine at the N-terminus (K3) is pink and the catalytic site is purple. Quantified data are representative of three independent experiments unless otherwise stated. Statistical significance was determined using one-way ANOVA.

USP18 has been reported as a positive regulator of cell growth in INF-1 treated cancer cells (Diao et al., 2020). Here, we investigated whether the two USP18 isoforms exert an effect on cell growth in muscle cells without INF-1 treatment. The doubling time of SF-expressing cells was approximately 20% shorter than that of LF-expressing cells (**Figure 1G**). Accordingly, LF-expressing cells proliferated significantly more slowly than SF-expressing or parental cells, whereas the proliferation rate of SF-expressing cells was comparable to that of parental cells (**Figure S1A**). These findings indicate that LF, but not SF, suppresses cell proliferation. Despite the difference in proliferation rate, the morphology of the two cell lines was indistinguishable (**Figure S1B**).

Next, we examined whether the two isoforms differentially regulate myogenic differentiation by quantifying MYOG expression. Western blot analysis showed that MYOG levels increased at day 2 of differentiation and decreased by day 3 in both LF- and SF-expressing cultures (**Figure S2**). Immunofluorescence analysis confirmed MYOG temporal expression and further revealed that, at day 1 of differentiation, SF-expressing cultures exhibited significantly higher MYOG fluorescence intensity than LF-expressing cultures (**Figure 1H-I**). In addition, the area of multinucleated objects, indicating fused cells, was significantly larger in SF cultures after one day of differentiation (**Figure 1J**), indicating that SF accelerates the early phase of myogenic differentiation. Myotube maturation was assessed by measuring myosin heavy chain (MyHC) expression. MyHC was not detected in either LF- or SF-expressing myotubes, whereas clear expression was observed in the non-transfected parental cells (**Figure S3**). These findings suggest that the lack of MyHC was therefore not specific to either USP18 isoform.

Endogenous USP18 levels at days 2 and 3 of differentiation were substantially lower than those observed with transgene overexpression and followed a similar pattern to MYOG expression (**Figure S3**). We then further explored the relationship between nuclear USP18 expression and myogenic differentiation, quantifying MYOG expression specifically in USP18-positive cells under proliferating conditions and after one day of differentiation. (**Figure 1K**). Nuclear USP18 fluorescence intensity decreased during differentiation in both LF- and SF-expressing cells. However, the proportion of USP18-positive nuclei decreased by approximately 50% in LF cultures but increased by nearly 40% in SF cultures (**Figure 1L**), suggesting greater stability or nuclear retention of the SF isoform during early differentiation. Among USP18-positive cells, MYOG expression was higher in SF-expressing cultures under proliferating conditions and remained elevated after one day of differentiation. Moreover, the proportion of MYOG-positive nuclei increased from 1.4% to 24.7% in SF cultures, whereas it remained unchanged in LF cultures (**Figure 1L**). Together, these findings indicate distinct effects of the two USP18 isoforms, with LF suppressing cell proliferation and SF promoting the onset of myogenic differentiation.

Given the distinct localization and functional effects of LF and SF, we next asked whether structural differences could explain their divergent activities. Folding modelling predicted that both isoforms share an identical structure but differ in the conformation of their N-terminal region (**Figure 1M**). The N-terminal region in LF formed an α-helix that folds back onto the α-helical region of the STAT2-binding domain, while in SF a α-helix is not formed, and the N-terminal adopted an open conformation (**Figure S4**). Notably, the N-terminus in the LF contains a single degron element with a conserved MSK (Kumar et al., 2022) sequence (**Figure S4**) with a lysine at position 3 (**Figure 1M**). These predicted structural differences may influence isoform-specific protein interactions and provide further insight into the scaffolding functions and stability of USP18.

### The USP18 interactome is affected by its isoforms

To gain insight into the molecular basis of the functional differences between the LF and SF isoforms, we compared their protein interactomes in proliferating cells and after one day of differentiation. FLAG immunoprecipitation followed by mass spectrometry (MS) was used to identify proteins interacting with each isoform (**Figure 2A**). Lysates from LF- and SF-expressing cells were subjected to FLAG immunoprecipitation (IP), while matched control samples were processed in parallel without anti-FLAG antibody (beads-only). Beads-only samples were used to determine a non-specific background. Western blot analysis of input samples confirmed expression of both USP18 isoforms (**Figure 2B**). Both LF and SF were detected in the IP under proliferating and differentiating conditions, confirming efficient capture of the tagged proteins. The FLAG signal was not detected in the beads-only controls, demonstrating the success of the pull-down (**Figure 2B**).

**Figure 2.**
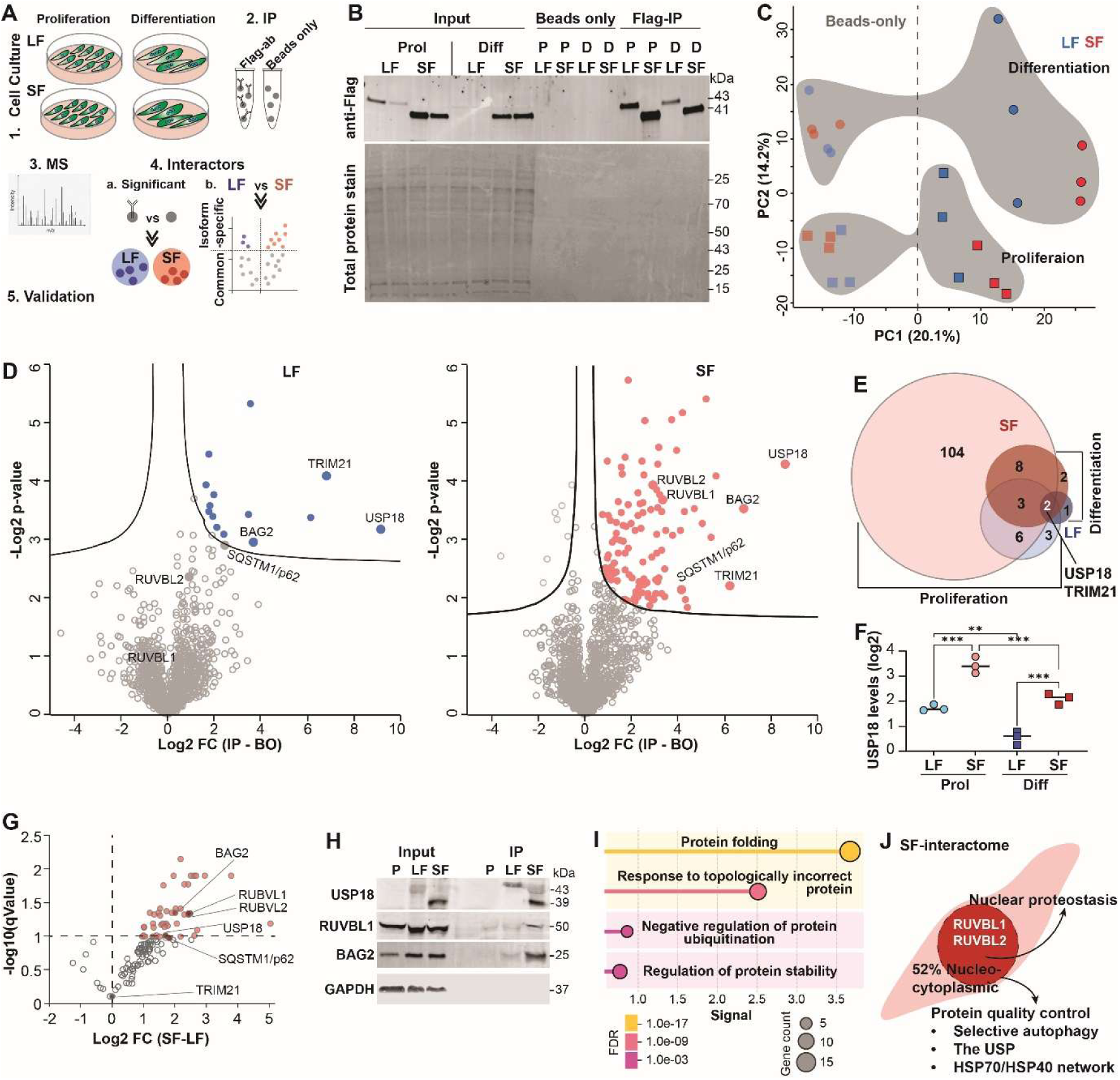
USP18 isoforms display distinct protein interaction networks during proliferation and early myogenic differentiation. **A.** Experimental workflow. Stable muscle cell lines expressing FLAG-tagged LF or SF were cultured under proliferating conditions or differentiated for 1 day. FLAG immunoprecipitation (IP) was performed together with beads-only controls, followed by mass spectrometry (MS), identification of significantly enriched interactors, and validation by immunoblotting. **B.** Representative Western blot of FLAG immunoprecipitation (Flag-IP) vs. Beads only control in proliferating (Prol) and differentiating (Diff) conditions. Input lysates confirmed LF and SF expression. Total protein staining served as a loading control. **C.** Principal component analysis (PCA) of the mass spectrometry dataset. PC1 (20.1%) separated FLAG-immunoprecipitated samples from beads-only controls, whereas PC2 (14.2%) separated proliferating (circle) and differentiating (square) samples. Gray area encircles samples from the same cell culture condition. **D.** Vulcano plots showing significant proteins in FLAG-IP relative to beads-only controls for LF (left) and SF (right). Significant interactors were identified using a 5% FDR are colored, and selected candidates, including USP18, TRIM21, BAG2, SQSTM1/p62, and RUVBL1/2, are highlighted. **E.** Venn diagram showing overlap of the significant interactors identified in proliferating and differentiating cells expressing LF or SF. TRIM21 and USP18 were detected in all four experimental conditions, whereas most interactors were isoform- and condition-specific. **F.** USP18 levels in proliferation (Prol) or 1 day of differentiation (Diff). USP18 abundance was normalized to the IgGs average in the mass spectrometry dataset. Data are presented as mean ± SE. Statistical significance was determined using two-way ANOVA, \*\**P* < 0.001. **G.** Vulcano plot of the significant Flag-IP proteins in LF vs. SF. Values normalized to IgG levels. SF-specific interactors (in red) were identified using a 5% FDR, and selected candidates, including USP18, TRIM21, BAG2, SQSTM1/p62, and RUVBL1/2, are highlighted. **H.** Validation of selected USP18 interactors by FLAG pull-down followed by Western blotting. GAPDH served as the loading control for the input samples. **I.** Enrichment analysis of shared (LF/SF) and SF-specific interactors. Shared interactors were significantly enriched for structural constituents of the cytoskeleton, whereas SF-specific interactors were enriched for protein folding, chaperone binding, chaperone-mediated autophagy, and sarcomeric proteins. Bubble size indicates the number of genes in each pathway, and color represents the false discovery rate (FDR). **J.** Schematic presentation of SF-specific nuclear interactors. The nuclear SF-specific interactors were clustered into four functional modules: nuclear proteostasis, genome maintenance, RNA metabolism, and nuclear organization.

We first assessed the source of variations between the 24 samples using the principal component analysis (PCA). The largest source of variation separated FLAG-immunoprecipitated samples from beads-only controls (PC1, 20.1%), indicating that the immunoprecipitated samples formed a distinct group from background proteins (**Figure 2C**). Proliferating and differentiating samples separated along PC2 (14.2%), suggesting that culture conditions also influence the isoform interactome (**Figure 2C**).

The significant interactors with each isoform were identified with a paired analysis against the beads-only samples (5% FDR). USP18 was found in all conditions, confirming successful immunoprecipitation (**Figure 2D** and **Figure S5**). Comparison of the significant interactors across the four conditions revealed that the size of the interactome differed markedly between isoforms and cellular states. SF exhibited approximately 11-fold more significant interactors than LF, while differentiation substantially reduced the number of interactors for both isoforms, with an 86% decrease for SF and a 33% decrease for LF (**Figure 2E**). These findings indicate that USP18 interactomes are dynamic during the early stages of myogenic differentiation. Approximately 80% of LF interactors overlapped with the SF significant interactors (**Figure 2E**) and significantly enriched cytoskeletal proteins (**Figure S6**). Tubulin proteins interacting with USP18 were previously reported (Jiao et al., 2017; Lin et al., 2026), suggesting a LF-SF common pathway. Across all conditions, only USP18 and TRIM21 were shared (**Figure 2E**). USP18 interaction with TRIM21 was also previously reported (Lin et al., 2026)., Normalized USP18 abundance was higher in SF than in LF samples and decreased upon differentiation for both isoforms, consistent with the immunoblot data (**Figure 2F**).

The isoform-specific interactors were then identified with a differential abundance analysis of the significant LF and SF interactors. We identified 47 SF-specific interactors, but none were LF-specific (**Figure 2G**). Amongst the 47 interactors were BAG2, p62 and RUVBL1/2, but TRIM21 was not SF-specific (**Figure 2G**). Co-immunoprecipitation experiments confirmed the stronger association between BAG2 and RUVBL1 with SF than with LF (**Figure 2H**). Functional enrichment analysis of the SF-specific interactors indicated pathways involved in protein folding, protein quality control, and proteostasis (**Figure 2I**). Consistent with the preferential nuclear localization of SF, more than half of these proteins were nuclear or shuttle between the nucleus and cytoplasm. Several interactors, including p62, members of the BAG protein family, and the ubiquitin-ribosomal fusion protein RPS27A/UBE2P80, were associated with protein quality control and nuclear proteostasis, whereas HSP40 and HSP70 family chaperones are key regulators of protein folding and stability (**Figure 2J**). In addition, SF interacted with the chromatin remodeling factors RUVBL1 and RUVBL2, suggesting that nuclear SF may also influence transcriptional regulation and genome maintenance. Together, these findings indicate that in proliferation conditions, the SF isoform preferentially associated with proteins involved in nuclear proteostasis and protein homeostasis.

### USP18-SF exhibits increased protein stability and altered proteostasis

Given the enrichment of autophagy- and protein quality control-associated proteins in the SF interactome, we next examined whether the two USP18 isoforms differentially affect components of the autophagy pathway. The SF-specific interactors are p62/SQSTM1 and BAG2. While p62 puncta are classic hallmarks of autophagic machinery (Bjørkøy et al., 2005), BAG2 forms a distinct type of puncta associated with stress response (Carrettiero et al., 2022). We quantified cytosolic p62 and BAG2 puncta (**Figure 3A**). SF-expressing cells displayed a significantly greater number of p62 and BAG2 puncta than LF-expressing cells (**Figure 3A**) consistent with altered p62 organization/protein-homeostasis pathways in SF-expressing cells.

**Figure 3.**
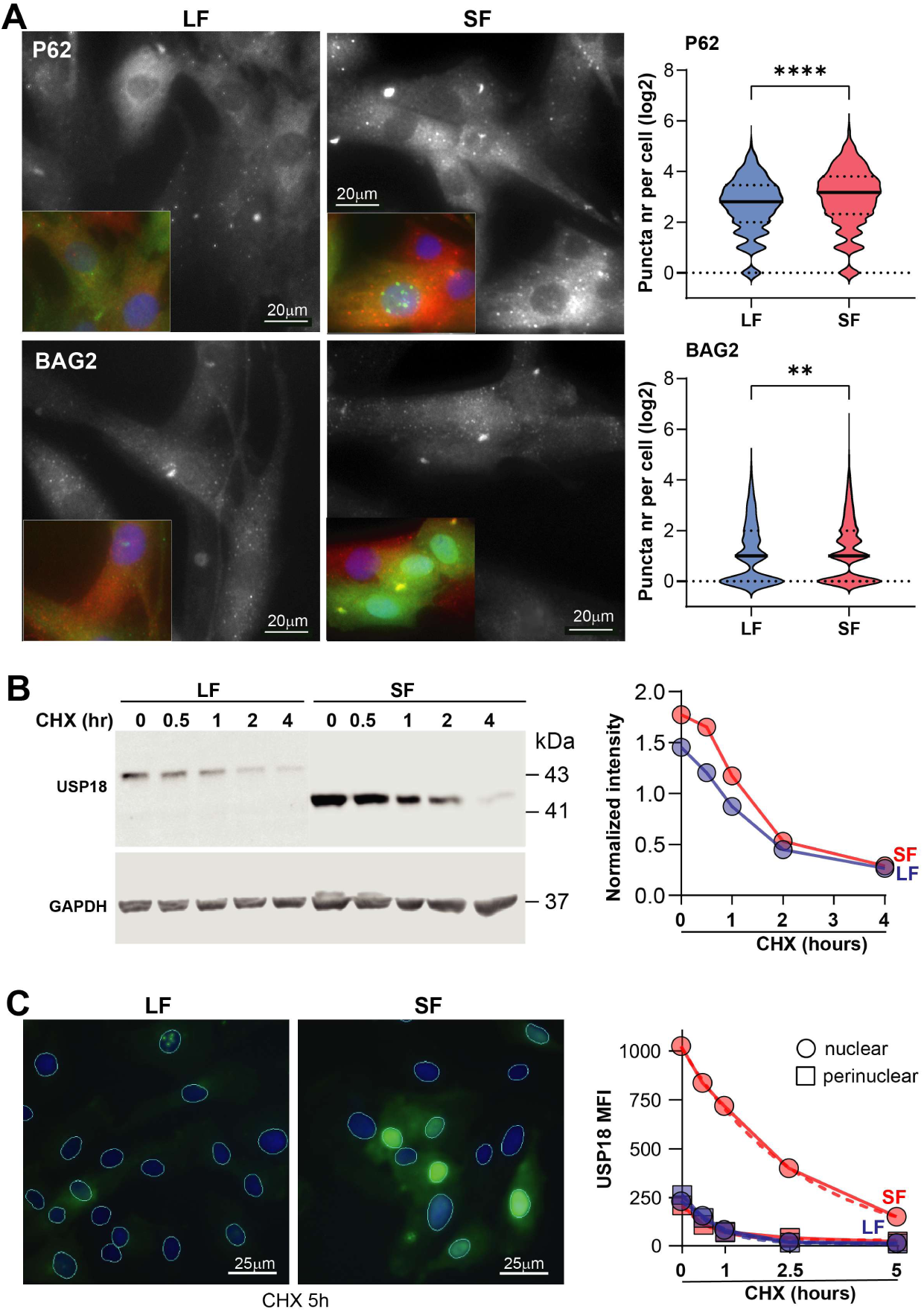
USP18-SF exhibits increased stability and nuclear accumulation and alters protein homeostasis-associated puncta. **A.** Representative images showing p62 and BAG2 puncta in USP18-LF- and USP18-SF-expressing cells. Insets show representative enlarged regions with nuclear staining. Right, quantification of p62 and BAG2 puncta per cell. SF-expressing cells displayed increased numbers of p62 and BAG2 puncta compared with LF-expressing cells. Statistical significance is indicated by asterisks (**P < 0.01, ****P < 0.0001), calculated from >1000 cells using Welch’s T-test. **B.** Representative immunoblot showing USP18-LF and USP18-SF protein levels following cycloheximide (CHX) treatment at the indicated time points. GAPDH was used as a loading control. Exposure is adjusted per isoform. Right, quantification of USP18 protein abundance following CHX treatment, normalized to the respective untreated condition. Hours indicate incubation with Cycloheximide. **C.** Representative fluorescence images of USP18-LF- and USP18-SF-expressing cells following 5 h of CHX treatment. USP18 signal was quantified separately in the nucleus and perinuclear region. USP18-SF showed increased retention in the nucleus following CHX treatment compared with USP18-LF.

The differential accumulation of the two isoforms under normal conditions, together with the enrichment of scaffold and proteostasis-associated proteins in the SF interactome, suggests that the two isoforms differ in protein stability. To test this, we assessed protein turnover following cycloheximide (CHX) chase treatment in cells by Western blot analysis. LF abundance decreased already after 30 minutes of CHX treatment, whereas SF abundance remained relatively stable and decreased only after 60 minutes, indicating that SF exhibited slower turnover/increased stability (**Figure 3B**). We then validated these findings by quantifying the FLAG signal in the nuclear and perinuclear compartments using immunofluorescence. in the perinuclear compartment, LF and SF showed comparable stability (**Figure 3C**). But in the nuclear compartment, SF stability was higher in both proliferation and differentiation conditions (**Figure S7**), raising the hypothesis that the SF plays a role in nuclear activities.

### Alternative splicing shapes distinct RNA expression profiles in LF and SF cell cultures

To determine whether the nuclear SF affects expression patterns during early myogenesis, we performed RNA sequencing on cells one day into differentiation. Differential expression analysis identified 1,393 genes distinguishing LF-from SF-expressing cells, with roughly 45% upregulated in SF (**Figure 4A**), indicating the more genes are upregulated by LF expression compared to SF. Notably, USP18 itself emerged as the single most upregulated gene in SF-expressing cells (**Figure 4A**). This USP18 upregulation represents the endogenous expression, since the transgene mRNA levels are similar between LF and SF (**Figure 1B**). This raises the possibility that SF positively regulates USP18 gene expression. Comparing our differentially expressed genes against a published USP18 knockdown dataset (**Figure S8**), we found that SF-upregulated genes overlapped significantly more with USP18-regulated genes than LF-upregulated genes did, indicating that USP18’s transcriptional effects during early myogenic differentiation were more strongly associated with the SF expression profile. Despite USP18’s canonical role in type-I interferon signaling, interferon-stimulated genes were not enriched among SF-upregulated genes, suggesting the transcriptional changes we observed reflect non-canonical SF-USP18 functions rather than classical interferon pathway activity.

**Figure 4.**
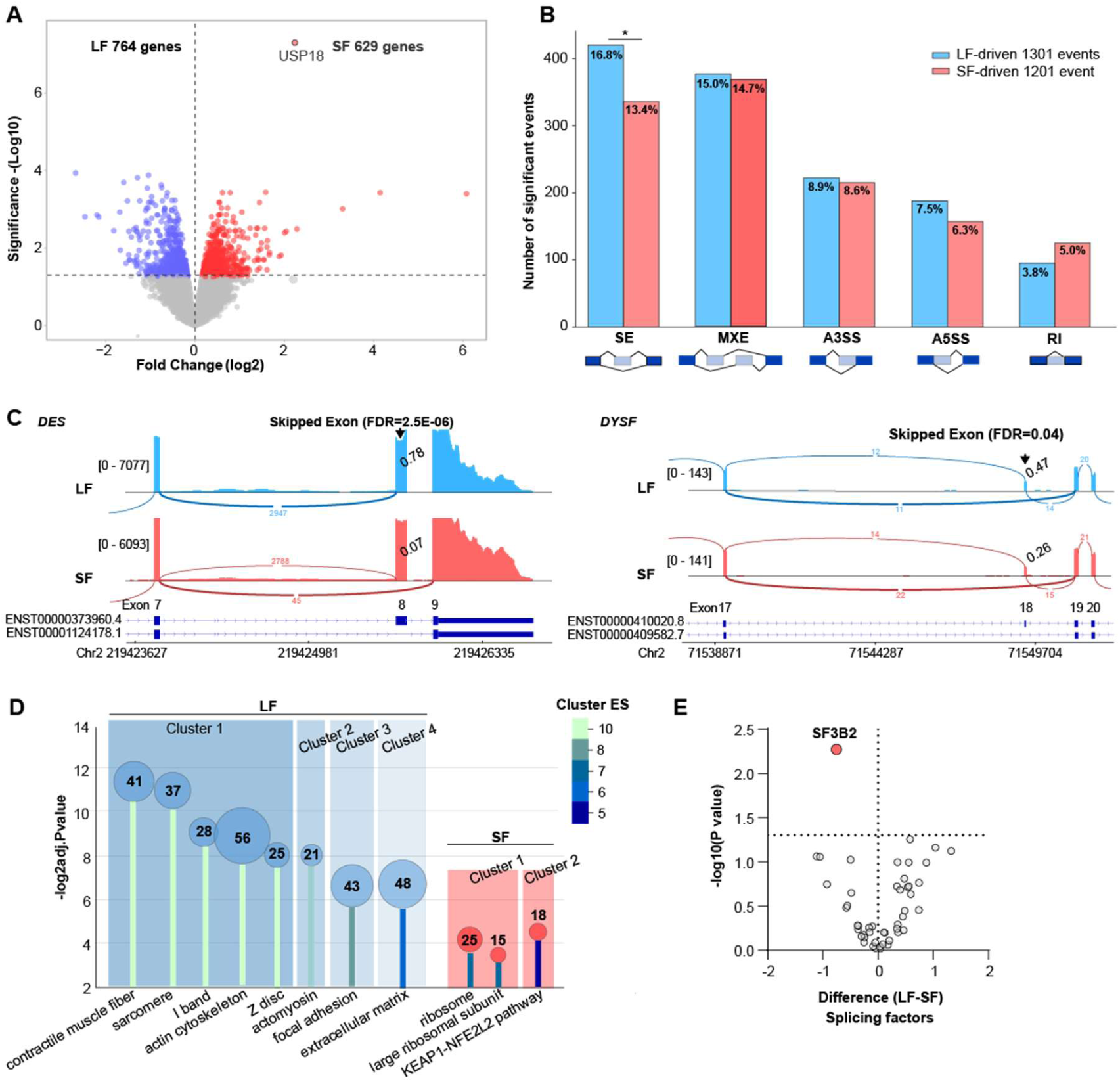
Differential gene expression and alternative splicing analyses between LF and SF USP18 isoforms. RNAseq was conducted in RNA from cell cultures grown in differentiation conditions for one day (N=3 per group). **A.** Volcano plot shows differential gene expression (log₂ fold change vs. -log₁₀ *P* value). The horizontal dashed line marks the significance threshold (FDR = 0.05). Genes significantly expressed in LF are blue and in SF are red; non-significant genes are grey. *USP18* is labeled. The number of DE genes per group is denoted. **B.** Bar chart of five alternative splicing events shows the number of significant events (FDR < 0.05 and |ΔPSI| ≥ 0.1) per event class: skipped exon (SE), mutually exclusive exons (MXE), alternative 5′ splice site (A5SS), alternative 3′ splice site (A3SS) and retained intron (RI). A schematic representation of each event class is shown below the axis: blue boxes, constitutive exons; white boxes, alternative exon (e*) or retained intron. Events with higher exon inclusion in LF (ΔPSI ≥ +0.1; n = 1301) are denoted in blue, and events with higher inclusion in SF (ΔPSI ≤ −0.1; n = 1201) in red. The proportion of events assigned to each group, relative to the 2,502 significant events, is indicated above each bar. The asterisk marks a significant deviation from an equal LF:SF split (binomial test against 50:50, Benjamini-Hochberg corrected across the five event classes; *P*_BH_ *<* 0.05). **C.** Sashimi plot examples of two skipped-exon events in muscle genes. Reads and exon skipping events in LF are in blue and SF in red. An arrow marks the significant SE event. Upper panel: *DES*, skipping of exon 8 (arrow; FDR = 2.5 × 10^-6^), PSI = 0.78 in LF and 0.07 in SF. Lower panel: *DYSF*, skipping of exon 18 (arrow; FDR = 0.04), PSI = 0.47 in LF and 0.26 in SF. Read coverage is shown as a histogram with the data range in brackets; arcs represent exon-exon junctions and the numbers on the arcs are the junction read counts. Ensembl transcript schematic view and exon numbers are shown below each track. **D.** Functional annotation clustering enrichment (DAVID) of the LF-DE (blue) and SF-DE (red) gene sets. The y-axis shows the significance of each annotation term (-log_2_ adjusted *P* value). The bubble area is proportional to the number of genes in the term. The color of the lollipop stem indicates the enrichment score (ES) of the annotation cluster to which the term belongs. Clusters are denoted with a colored box. **E.** Volcano plot of splicing factors. Values normalized to IgG levels were used for differential abundance analysis. Significance was assessed by unpaired t-test comparing LF and SF interactors across proliferation and differentiation conditions combined.

Given that the SF interactome was enriched for nuclear RNA-metabolism proteins, we next investigated whether the transcriptomic differences extended beyond changes in gene expression by quantifying alternative splicing events across the five major classes: skipped exons (SE), mutually exclusive exons (MXE), alternative 3′ splice sites (A3SS), alternative 5′ splice sites (A5SS), and retained introns (RI) (Li et al., 2026). We identified 2,502 significant events overall (FDR < 0.05, |ΔPSI| ≥ 0.1). Of these classes, only skipped exon events differed significantly between the two isoforms (Benjamini-Hochberg-adjusted P = 0.013) (**Figure 4B**). A higher number of ES events in LF points to reduced exon skipping in SF-expressing cells. To validate these computational findings, we visualized splicing events in two muscle-associated genes (**Figure 4C**). *DES* (desmin), which anchors the sarcomere to the sarcolemma, showed markedly reduced inclusion of exon 8 in SF cells (PSI 0.78 vs. 0.07; FDR = 2.5 × 10⁻⁶). *DYSF* (dysferlin), essential for membrane repair, showed a similar though smaller reduction in exon 18 inclusion (PSI 0.47 vs. 0.26; FDR = 0.04).

Pathway enrichment analysis of both differentially expressed and alternatively spliced genes helped clarify the biological processes involved. Genes upregulated in LF were enriched primarily for muscle organization pathways, while genes upregulated in SF showed fewer and weaker enrichments overall (**Figure 4D**). By contrast, the full set of 1,620 alternatively spliced genes, spanning all five event classes, was significantly enriched for nuclear processes, DNA damage response, transcriptional regulation, and RNA processing (**Figure S9**). This pattern aligns with SF’s nuclear localization and its preferential binding to RNA-metabolism and nuclear-homeostasis proteins. Although myogenic pathways themselves were not significantly enriched, several key regulators of muscle differentiation were nonetheless subject to alternative splicing, suggesting that coordinated splicing changes, rather than pathway-wide shifts, may underlie the phenotypic differences observed.

As SF may affect transcriptional systems, we assessed whether there are splicing factors interacting differently between the two isoforms. 40 splicing factors were found in the dataset, amongst which only SF3B2 significantly interacted with the SF (**Figure 4E**). SF3B2 has been implicated in muscle function (Kitamura et al., 2022; Murphy et al., 2026).

## Discussion

USP18 is best known for its dual role in IFN-I signaling, acting both as an ISG15 deISGylase and as a non-catalytic scaffold regulating IFNAR signaling and it also has reported deubiquitinase activity (Kang & Jeon, 2020). Through these activities, USP18 is an important regulator of immune responses, with implications for antiviral host defense and cancer progression. Although USP18 expression is low in most cultured cells, it can be induced by stress and inflammatory signals independently of IFNs, suggesting additional functions. (Kang & Jeon, 2020). In muscle cells, USP18 plays a non-canonical role at its base-level, suggesting a role in muscle regeneration and differentiation (Nijssen et al., 2026; Olie et al., 2023).

In the present study, we describe isoform-specific functions of USP18 in muscle cells. In particular, the SF isoform enhanced of myogenic differentiation at an early step, whereas the LF isoform promoted cell growth arrest. Most strikingly, we show that the SF interactome is substantially larger than the LF’s, suggesting a structural role of the N-terminal in protein interactions that grants USP18-SF novel cellular functions. The difference in the N-termini of the LF and SF isoforms places USP18 amongst a subtype of N-terminus proteoforms, defined as protein variants with altered N-termini generated through RNA-driven processes (Morrison & Rissland, 2025). In the case of USP18, these proteoforms arise from differential translation of the initiation site usage. N-terminal sequences can influence protein stability, for example through the presence of E3 ligase recognition motifs that promote protein degradation (Morrison & Rissland, 2025).

The presence of MSK at the N-terminus of the LF could align with shorter stability compared to the SF. The full length of USP18 functions as a scaffold affecting TRIM21-TRIM29 E3-ligase complex (Lin et al., 2026), suggesting that USP18 affects ubiquitination through its scaffold function.

Several SF-specific interactors were also associated with protein stability and cellular stress, including BAG2 and p62. We observed a higher number of BAG2 and p62 puncta in SF-expressing cells suggesting a stronger effect of the SF isoform on autophagy- and protein homeostasis-related pathways. p62 puncta indicate autophagic activity or protein turnover, while BAG2 puncta indicate increased proteostatic or cellular stress (Carrettiero et al., 2022). We recognize that high overexpression levels can affect the protein degradation machinery, even when the overexpressed protein is not intrinsically proteotoxic (Moriya, 2015). Therefore, the increased BAG2 and p62 puncta observed in SF-expressing cells could reflect an overexpression effect.

A prominent feature of the USP18 SF interactome is the large number of nuclear proteins. The majority shuttle between the cytoplasm and nucleus, including stress-related proteins such as BAG2 and heat-shock proteins, whereas a smaller subset predominantly resides in the nucleus. Among the latter are RUVBL1 and RUVBL2, which function as transcriptional co-activators and form a complex with RNA polymerase II (Pol II)(Qiu et al., 1998) thereby influencing genome-wide RNA expression profiles. Here, we show that LF and SF display markedly different gene expression profiles. More strikingly, however, we observed a significant reduction in exon-skipping events in the SF transcriptome. We identified the splicing factor SF3B2 as a preferential interactor of SF compared with LF. SF3B2 also interacts with Pol II and modulates its activity (Kitamura et al., 2022). Together, these findings raise the possibility that a complex containing Pol II, SF3B2, RUVBL1, RUVBL2, and USP18-SF contributes to post-transcriptional regulation during myogenesis, with SF binding to SF3B2 potentially suppressing its activity. Interestingly, among the genes showing exon-skipping events are *desmin* and *dysferlin*, both encode proteins important for skeletal muscle structure or function. Thus, SF may regulate myogenesis, in part, through modulation of alternative splicing, a process known to play an important role during myogenesis (Donega et al., 2025). Our findings expand the functional landscape of USP18 beyond its established role in IFN-I signaling and suggest that USP18 isoforms may provide an additional layer of regulation. We suggest that the short isoform of USP18 has a nuclear function that affects alternative splicing and may contribute to an early step of myogenesis and propose that USP18 represents an additional example of an N-terminal proteoform in which N-terminal variation influences protein stability, protein–protein interactions, and subcellular localization.

## Acknowlegements

We thank members of the Raz, Pinto-Fernández and Kessler groups for insightful discussions related to the manuscript.

## Author Contributions

T.I.N and S.D.D. performed biochemical experiments. T.I.N. and M.S. performed cell biology experiments. M.S. performed RNAseq data analysis. S.F. and I.V. performed mass spectrometry and proteomics analysis. A.P.F. supervised biochemical experiments. A.P.F., B.M.K. and V.R. conceptualised the study. V.R and T.I.N. wrote the manuscript, and all authors commented and contributed to editing the paper.

## Data Availability

MS data has been uploaded to the PRIDE data repository and is available to download under the accession number PXDxxxxxx (Perez-Riverol et al., 2025). RNAseq data has been uploaded to the EGA data repository and is available for download under the accession number EGAD50000003142.

